# Transcriptional differences for COVID-19 Disease Map genes between males and females indicate a different basal immunophenotype relevant to the disease

**DOI:** 10.1101/2020.09.30.321059

**Authors:** Tianyuan Liu, Leandro Balzano Nogueira, Ana Lleo, Ana Conesa

## Abstract

Worldwide COVID-19 epidemiology data indicate clear differences in disease incidence among sex and age groups. Specifically, male patients are at a higher death risk than females. However, whether this difference is the consequence of a pre-existing sex-bias in immune genes or a differential response to the virus has not been studied yet. We created DeCovid, an R shiny app that combines gene expression data of different human tissue from the Genotype-Tissue Expression (GTEx) project and the COVID-19 Disease Map gene collection to explore basal gene expression differences across healthy demographic groups. We used this app to study differential gene expression between men and women for COVID-19 associated genes. We identified that healthy women present higher levels in the expression of interferon genes and the JAK-STAT pathway leading to cell survival.

---

On the contrary healthy men exhibit higher levels of pro-inflammatory cytokines. These pathways have been associated with the severity of COVID-19 progression. Our data suggest that men and women have a different basal immunophenotype that is relevant to the severity of COVID-19 and may explain differences in fatality rates between the two gender groups. The DeCovid app is effective and easy to use tool to explore the expression levels of genes important for SARS-CoV-2 infection among demographic groups and tissues. The DeCovid app is publicly available at https://github.com/ConesaLab/DeCovid.

## 1 Introduction

The severe acute respiratory syndrome coronavirus 2 (SARS-CoV-2), the virus that causes coronavirus disease 2019 (COVID-19), is without doubt the most severe health, economic and social threat of our time. The COVID-19 has followed two infection major waves in Spring and Fall 2020 and have reached the one million deceased people by end September 2020. The understanding of the disease has evolved over time. While initially COVID-19 was considered as severe atypical pneumonia, it soon became clear that the virus can infect many tissues. Moreover, the disease affects blood coagulation and can cause an exacerbated immune response referred to as the “cytokine storm”. Although clinical management of COVID-19 has rapidly evolved over time, no curative treatment is yet available, and current recommendations support the use of steroids, oxygen, and a prophylactic dose of heparin. While accelerated vaccine development efforts have resulted in tremendous advances in a relatively short time, the WHO acknowledges that a vaccine solution for the general population might not be ready before mid or late 2021. At the same time, very recent re-infection cases suggest that immunity may not always persist, and active disease surveillance of past cases may be required. As the second wave of the pandemic steadily progresses in Europe and the US, possibly with new variants of the virus and affecting a more diverse and younger population, novel or improved treatment targets and options are likely to remain as important for the management of the disease.

One of the most intriguing aspects of COVID-19 is the different grade of severity it affects different people. Although the acknowledged risk factors such as pre-existing conditions such as cardiovascular disease (Zheng *et al*., 2020; D. Wang *et al*., 2020; Chen *et al*., 2020), diabetes (Apicella *et al*., 2020; Fang *et al*., 2020; Guo *et al*., 2020, 19), obesity (Simonnet *et al*., 2020), age (Liu *et al*., 2020; CDC COVID-19 Response Team, 2020; LitCovid) and sex (higher mortality in men than women (Gebhard *et al*., 2020; Jin *et al*., 2020) suggest a relationship between the physical condition and disease progression, the precise pathways of this relationship have not been clearly established yet. Sex-associated differences between males and females that have been proposed to affect COVID-19 incidence include lifestyle (e.g., smoking and drinking) and mental health. The smoking population of males is higher than females, and smoking may be a risk factor for severe COVID-19 (Cai, 2020, 19; Jamal *et al*., 2016; Grundy *et al*., 2020). In contrast, men differ from women during the pandemic in sleeping and signs of depression (C. Wang *et al*., 2020). A recent study showed that male and female COVID-19 patients differed in their immune response, with the first showing a stronger cytokine response while the second is having a higher T-cell activation pattern (Takahashi *et al*., 2020). Interestingly, the pattern of COVID-19 risk factors is not fully shared with other similar recent pandemics such as SARS, MERS, H1N1. For example, SARS patients were more frequent among the healthy young people (Yin and Wunderink, 2018). Age and complications, but not gender, was the most significant risk factors for mortality in the Arab and South Korean Studies of MERS (Park et al., 2018). Several studies on the 2009 H1N1 influenza revealed the younger age, chronic conditions, and female sex as risk factors of the disease (Viasus *et al*., 2011). This suggests that the observed COVID-19 risk factors, especially those associated with age and sex groups, might be the result of specific interactions of the SARS-CoV-2 with intrinsic physiological characteristics, including the basal immunophenotype, of these population groups. However, which specific immunological characteristics imprint an existing condition that how this relates to COVID-19 remains to be explored.

The COVID-19 Disease Map initiative was launched in May 2020 with the aim of creating a collection of genes and pathways relevant to SARS-CoV-2 infection derived from the growing COVID-19 literature. This resource contains over 1,500 COVID-19 disease genes distributed among 221 pathways (Ostaszewski et al., 2020), including cytokine and interferon genes, and life cycle of SARS-CoV and MERS-CoV pathway. The COVID-19 Disease Map is a valuable resource to investigate the molecular responses to SARS2-CoV-2 infection and understand the biological pathways leading to the severe manifestations of the disease. This database also creates an opportunity for interrogating existing molecular data on differences associated with COVID-19 risk factors in the general population.

In this paper, we present the DeCovid app, a Shiny app, to explore basal expression level differences in COVID-19 disease map genes between men and women and different age groups. We used data from the GTEx database, which contains RNA-seq profiles for hundreds of demographically diverse healthy individuals in multiple tissues and allows us to interrogate COVID-19 genes globally or individually. We used this resource to study basal expression differences in COVID-19 genes between man and woman and found that a different immunological state is prevalent in both genders. Interestingly some of these differential pathways have been shown to be relevant for disease severity progression and be characteristic of the sex-biased immune response to the virus, providing ground for hypotheses on the molecular basis of the COVID-19 sex-bias. We anticipate that the DeCovid app could be a useful tool for researchers to explore the molecular etiology of COVID-19 demographic differences.

## Results

### DeCovid as a user-friendly tool to explore gene expression of COVID-19 genes in healthy individuals

The DeCovid shiny app combines a selection of human tissue specific GTEx data with the COVID-19 Disease Map database to allow quick exploration of basal gene expression values and differences in the healthy human population for genes described to be important for COVID-19. We included data from blood, lungs, heart, kidney, stomach, and brain for being tissues described to be affected by SARS-CoV-2 infection. The sample size for different tissues is different in the GTEx datasets. The largest sample size is the whole blood with 330 samples, while the smallest dataset is the stomach tissue with 57 samples (**Figure 1a**). The total number of collected COVID-19 Disease Map is genes 1,523. A Gene Ontology (GO) term enrichment analysis indicates that genes related to cytokines, response to virus, innate immune response, transcriptional regulation, and kinase activity populate this collection (**Figure 1b**).

**Figure 1.**
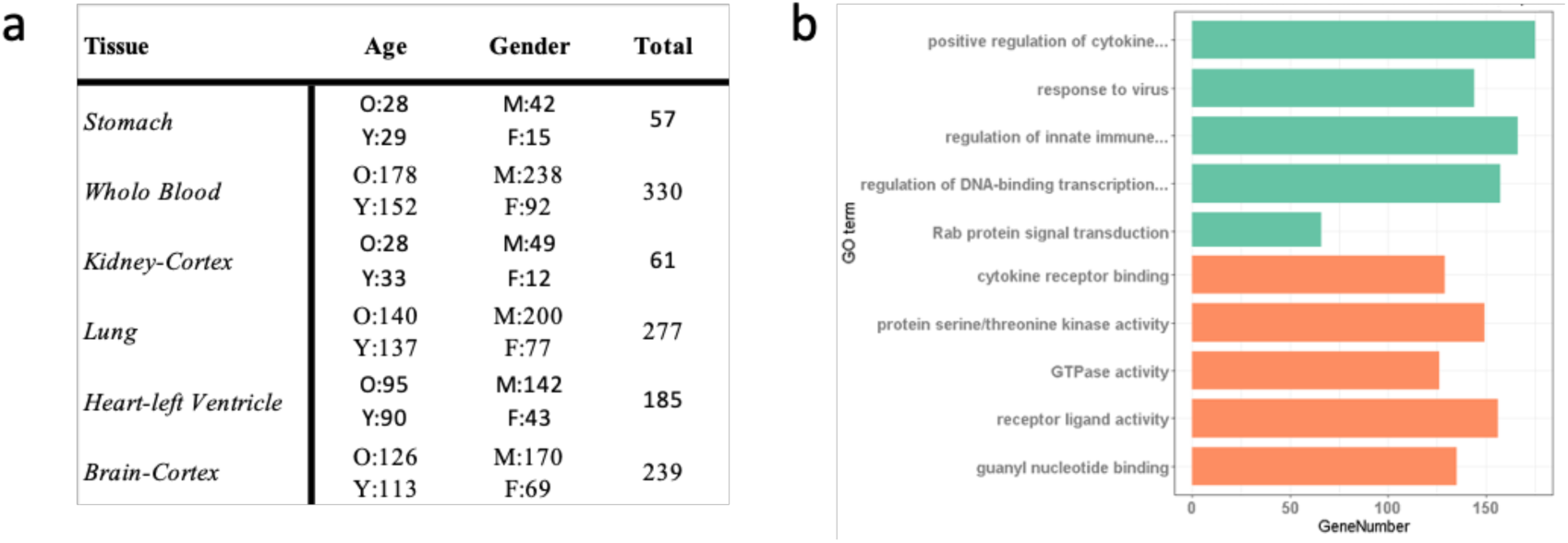
**a** Summary of the GTEx samples implemented in the DeCovid app. O: old age (age ≥ 60); Y: young age (age < 60); M: males, F: females. **b** Gene Ontology enrichment analysis of the 1,523 COVID-19 Disease Map genes. The most enriched GO term by category (green: Biological Process, orange: Molecular function) are shown.

The DeCovid app is a user-friendly implementation to allow an analysis of these data for researchers without strong bioinformatics skills (**Figure 2**). Extensive documentation and video tutorials are provided to the facility for a quick start. Users should indicate the demographic factor for differential expression analysis. In case age is selected, the age threshold value to classify samples as old or young should be specified. Once a significance *p*.*value* and a fold-change threshold are provided, the app computes differential expression and provides results as a list of genes and heatmaps with sex and age groups mean values (**Figure 2**). Additionally, users can explore the distribution of gene expression values for specific genes of the COVID-19 Disease Map collection or investigate the enrichment of specific immune response-related functions among genes with sex or age expression biases.

**Figure 2.**
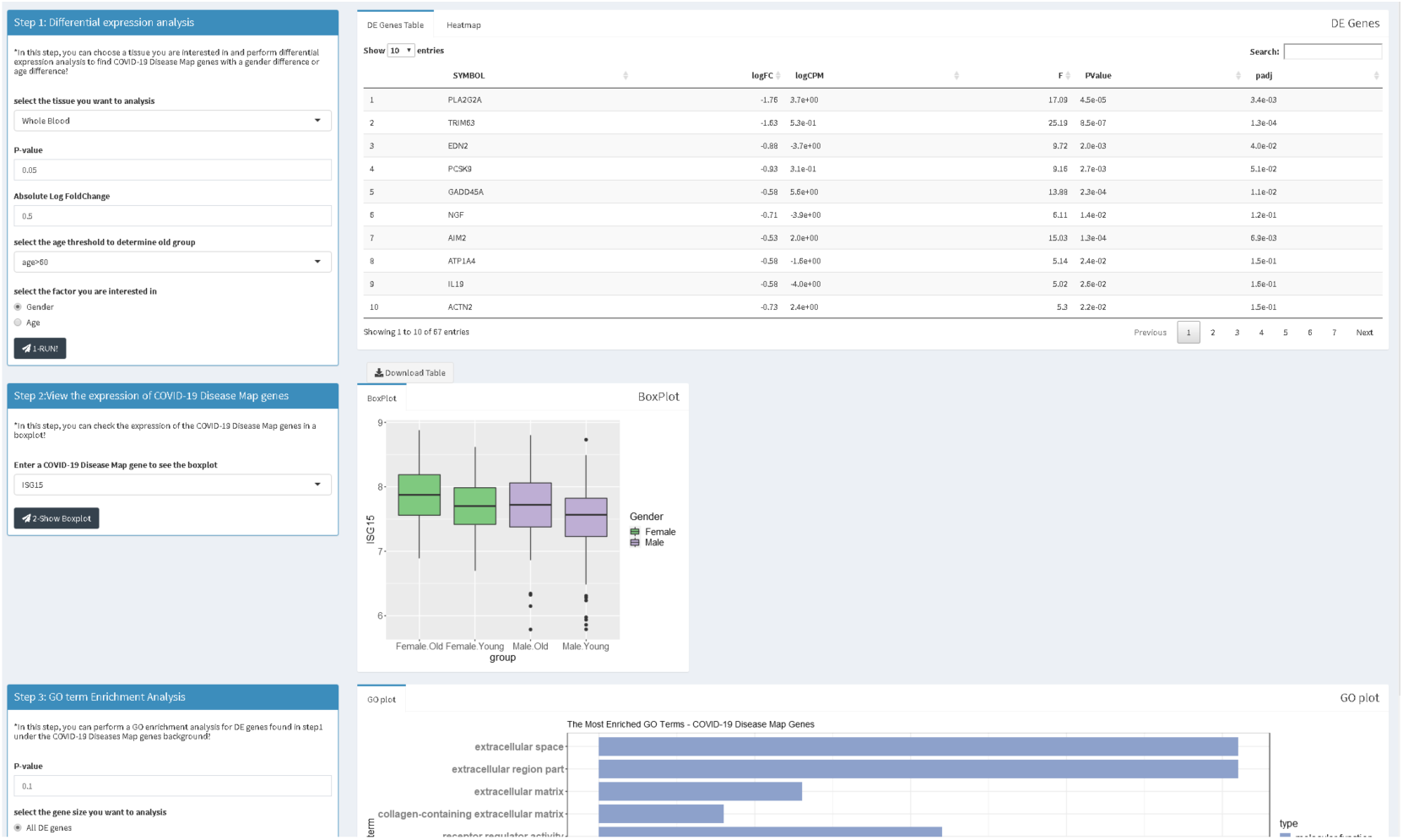
Screenshot of the DeCovid app showing on the left dialog panels for input parameters (contrast type and significance thresholds), and on the right results panels with list of differentially expressed genes, gene-specific expression plots and enrichment analysis.

### DeCovid shows wide-spread gene expression differences in COVID-19 Disease Map genes between men and women and age groups at multiple tissues

Using DeCovid, we analyzed the extent of gene expression differences among demographic groups across different COVID-19 affected tissues (Table 1). The analysis revealed a wide-range of significant gene expression differences for COVID-19 genes between when comparing sexes and age groups for all evaluated tissues. Differences where lower when imposing a fold-change threshold of 0.5 for the magnitude of the mean expression level. Whole-blood were the tissue with the highest number of expression differences for COVID-19 genes followed by the brain. From this simple analysis, we concluded that the COVID-19 Disease Map represents a disease signature with significant differences across demographic groups that may be worth exploring for hypothesizing on the molecular basis for differences in disease incidence among these groups.

**Table 1.**
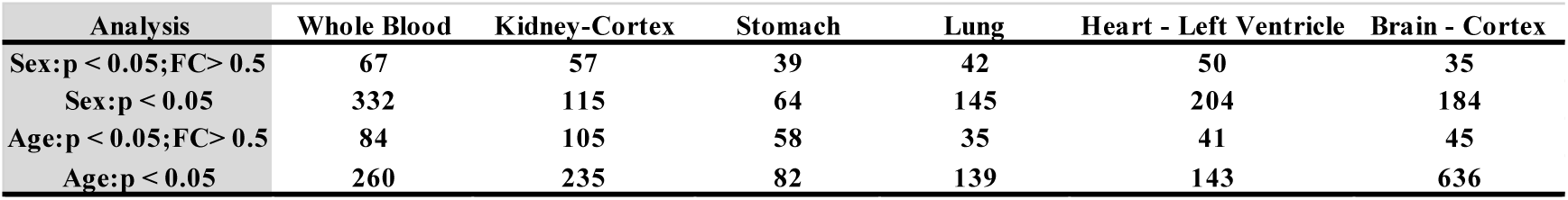
Number of differentially expressed COVID19 disease map genes between sex and age groups in different human tissues. Number of significant (p.value (p) < 0.05) genes are reported either applying or not a fold-change (FC) cut-off value of 0.5

### DeCovid identified basal sex-biased expression differences in immune-response genes associated to differential disease incidence between man and women

Since COVID-19 fatality rates are consistently higher in men than women in many different countries worldwide (**Supplementary Figure 1**), we hypothesized that sex is a pre-existing condition that influences the severity of the disease and that the study of basal differences in expression differences in COVID-19 relevant genes in healthy individuals could reveal insights into the molecular basis of these differences. We used the DeCovid app to explore these differences and concentrated on the blood tissue for capturing systemic immune responses. A total of 67 genes (p.value < 0.05 and fold change > 0.5) were differentially expressed between men and women in blood samples, regardless of the age group (**Figure 3a**). We found high expression in females of genes involved in the innate immune response, particularly type I interferons, such as IFNA17, IFNA2, IFNE, IFIT1, and IFIT3, anti-apoptotic regulators such as BCL2A, BCL2L1, and MCL1, and chemokines (CCL21, CXCL12) as well as the tumor necrosis factor (TNFSF13B). On the other hand, males showed higher basal expression proinflammatory chemokines (CXCL2) and cytokines (IL17, IL22, and IL3) genes. These gene expression differences point to two major immune response pathways differently active in men and women: interferon-mediated antiviral responses and the cytokine activation response. Interestingly these two processes have been indicated as distinctive pathways in the severity of the SARS-CoV-2 response (Hadjadj, J. et al. 2020).

**Figure 3.**
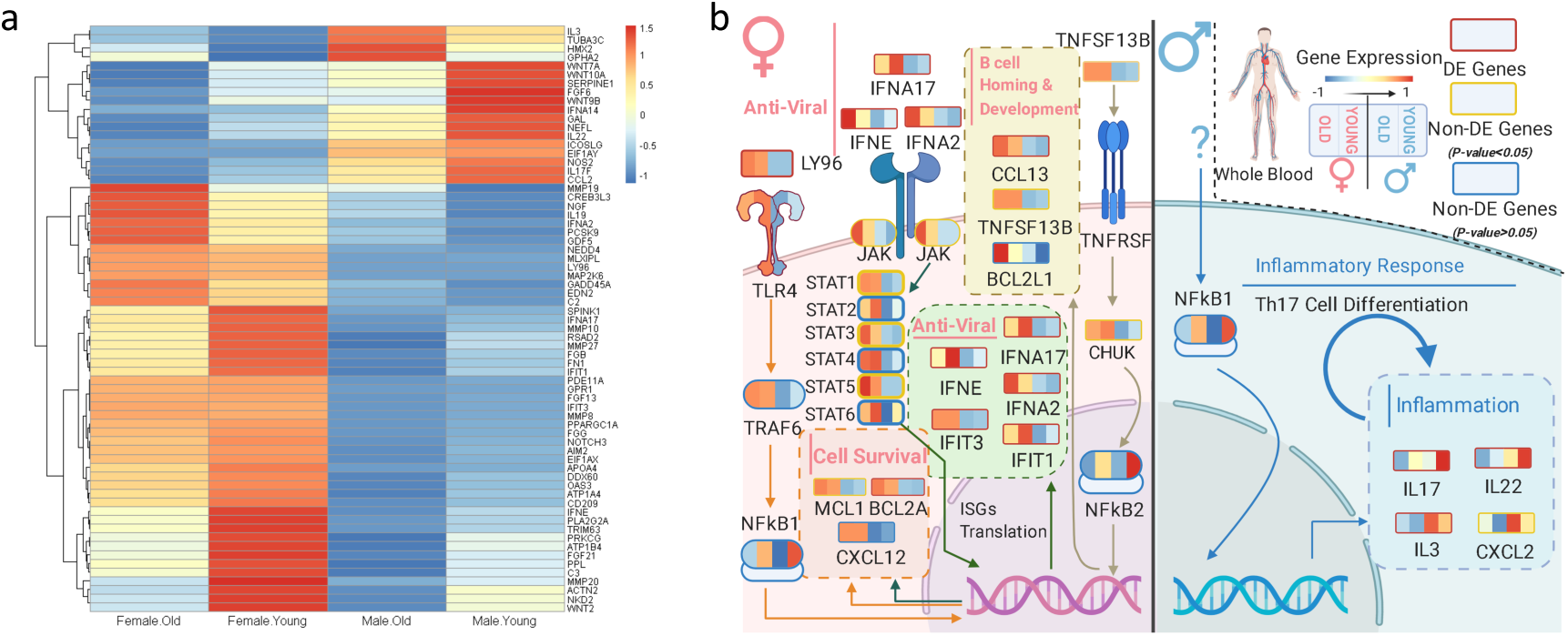
Analysis of COVID-19 Disease map gene expression differences between men and women in blood. **a** Heatmap of significant genes (p.value < 0.05 and FC > 0.5). Men expression values at each demographic group are shown. Old > 60 yrs; Young < 60 yrs. **b** Biological model for the differential gene expression. Small heatmaps on top of each gene name display consecutively mean expression values for healthy female_old, female_young, male_old and male_young groups. The model indicates that females present a more activated anti-viral response than males for anti-viral interferons, B-cell homing and cell survival pathways, while males have higher basal levels of pro-inflammatory cytokines (Created with biorender.com/).

Data suggest that females present a faster anti-viral response than males based on three main aspects. First, the higher levels in the blood of anti-viral type 1 interferons IFNA17, IFNA2, IFIT1, IFIT3, and IFNE, which serve as the first line of defense against viral and bacterial infections (Boxx and Cheng, 2016). Moreover, type 1 IFN enhances T-cell survival and effector functions through initial antigen recognition (Huber and Farrar, 2011), which may facilitate restricted virus proliferation. This transcriptional expression profile is in agreement with recent clinical studies that showed that female patients have a high T-cell level (Takahashi *et al*., 2020), which were postulated as a critical factor for the differential fatality incidence between sexes. Therefore, with a high level of baseline expression of type 1 interferons, female patients are more likely to maintain a high T-cell level in the early stage, which might lead to a milder evolution of the disease.

The relatively high levels of type 1 IFN in females could be related to the higher expression of genes in the TLR4 and TRAF6 genes, which leads to the synthesis of these cytokines (Makris *et al*., 2017). The TLR4/TRAF6/ NFkB1 pathway is likely more activated in females, possibly due to the upregulation of the gene LY96 (lymphocyte antigen 96), which stimulates activation of toll-like receptor 4 (TLR4) cascade leading to the synthesis of type 1 IFN (Makris *et al*., 2017). LY96 significant upregulation, as well as higher levels of TLR4, has been associated with resistance against viral infections in several species (Shinya *et al*., 2012), including mice models where mutations at TLR4 (Khanolkar *et al*., 2009) and type I IFN receptor (Haag *et al*., 2009) were shown to be critical to control hepatitis virus strain 1 (MHV-1) infection. Here we speculate that these innate immunity pathways may play a crucial role in the early control of respiratory coronavirus infection (**Figure 3b**). Moreover, data indicate higher expression in women of JAK, multiple members of the STAT family, and cell survival genes such as MCL1 and BCL2A. It has been shown that the Jak-STAT signaling pathway mediates the synthesis of anti-apoptotic regulators such as MCL1, BCL2A, and BCL2L1 upon induction of high type 1 interferon levels (Sepúlveda *et al*., 2007), agreeing with the observed gene expression phenotype in females.

Finally, we detected significant increased tumor necrosis factor TNFSF13B expression in females. This gene stimulates the synthesis of chemokines CCL21 and CXCL12 through NFkB subunit 2 pathway, leading to B-cell homing through the synthesis of CCL13 (López-Giral *et al*., 2004), lymphopoiesis through the synthesis of CXCL12 (Silva *et al*., 2016; Sen, 2006), and development and survival through the synthesis of TNFSF13B (Caamaño *et al*., 1998; Gerondakis and Siebenlist, 2010). Since B-cells are part of the humoral immunity component of the adaptive immune system and are responsible for mediating the production of antigen-specific immunoglobulins (Ig) directed against invasive pathogens, higher levels of these chemokines in females might lead to a faster response against a potential viral infection. In males, in turn, the mentioned processes were comparatively downregulated, while the only process upregulated compared to females were those related to the expression of pro-inflammatory cytokines (**Figure 3b**), particularly IL17, IL22 and IL3. Also, chemokine CXCL2 levels are higher in men, which supports the hypothesis of a first stage of male response against a viral infection being more related to inflammation and Th17 cell differentiation processes.

## Discussion

Here we present the DeCovid app as a resource to explore sex and age differential expression patterns in the healthy population for genes described to be involved in COVID-19 disease pathways. The GTEx data, used in this work has been recently mined for sex-specific expression differences concluding that these are tissue-specific and generally small, also observed in our analyses(Oliva *et al*., 2020). This study focused on the sex-specific genetic effects on gene expression associated to complex genetic traits. In our application we repurpose the valuable GTEx database for the study of COVID-19 relevant pathways while providing a mechanistic model to interpret differential expression results.

We used DeCovid to investigate the gene expression differences between men and woman that could explain differences in disease severity among these demographic groups, with men being more likely to have a fatal outcome than women. We found that two major immune response pathways were differentially expressed between men and women in the healthy population. Women showed higher expression of interferon genes while men have higher levels of pro-inflammatory Th17 cytokine genes. These two pathways have been proposed to be critical factors for fatal response to the virus infection, with patients that present an interferon-mediated response having a better prognosis than those responding with a massive cytokine activation (Blanco-Melo *et al*., 2020, 19; Hadjadj *et al*., 2020). In agreement, animal models of SARS-CoV-1 and MERS-CoV infection showed that failure to elicit an early type 1 interferon response correlates with the disease severity (Channappanavar *et al*., 2016). Perhaps more importantly, these models demonstrate that timing is key, as type 1 interferons are protective at the early stages of a viral disease. On the other hand, clinical studies showed a differences in the immunophenotype between female and male COVID-19 patients with women having a more robust T-cells activation while male patients presenting a higher plasma level of pro-inflammatory cytokines (Takahashi *et al*., 2020). In this study we showed that these immune system differences, which are critical to COVID-19 progression, represent a sex-associated pre-existing condition that is likely to create a differential predisposition to disease outcome between men and women and may explain the observed sex-bias in COVID-19 mortality. Future studies should address whether these gene expression patterns translate into functional differences in patients infected with SARS-CoV-2, whether they have prognosis value and of if they present any relationship with hormonal levels, which have also been linked to COVID-19 severity (Strope *et al*., 2020; Channappanavar *et al*., 2017; Scully *et al*., 2020). All together our results show that DeCovid is a useful resource to investigate the status of COVID-19 genes across demographic groups and human tissues and postulate on the differential pathways to disease operating in each case.

## Methods

### Datasets

We used RNA-seq data from the Genotype-Tissue Expression project (GTEx) that contains data from 44 tissue sources obtained from deceased but considered healthy individuals from both sexes and a wide range of ages(Lonsdale *et al*., 2013). Gene read counts matrix and the annotation files from the GTEx portal (https://www.gtexportal.org/home/datasets). In other to provide a homogenous set of human tissue reference samples, GTEx data were analyzed for possible biases, and samples belong to individuals with a “ventilator case” label as a cause of death were discarded as they clearly segregated from remaining samples on a Principal Component Analysis plot (**Supplementary Figure 1**). The COVID-19 Disease Map genes were downloaded from https://github.com/wikipathways/cord-19 and available on March 17th, 2020. The list is mined from 9996 PMC papers in COVID-19 Open Research Dataset (CORD-19) using machine learning approaches, so they are positively related to the COVID-19 disease process (Lu Wang et al., 2020).

### DeCovid software

The DeCovid software is a Shiny app written in R with a user-friendly interface. The app can be installed through a Docker image or directly download from GitHub (See **supplementary materials for installation**). DeCovid already has integrated all essential data from GTEx, and no additional downloads are necessary. Differential gene expression is calculated using edgeR (Robinson *et al*., 2010) and multiple testing correction is applied following the Benjamini and Hochberg(BH)method (Benjamini and Hochberg, 1995). Results are provided either as lists of differentially expressed genes, heatmaps of sex and age mean expression values, and gene-specific boxplots showing the distribution of expression values across demographic groups. GO enrichment analysis of significant gene sets is provided through the cluster profile R package and uses the list of COVID-19 disease map as a reference set (Yu *et al*., 2012).

## Authors contributions

TL: Created DeCovid app and performed statistical analyses

LB: Contributed in the interpretation of the DeCovid results on blood samples

AL: Supervised and contributed to interpretation of the DeCovid results on blood samples

AC: Conceived, coordinated and supervised the study.

## Notes

### Competing Interest Statement

The authors have declared no competing interest.

