## Supplementary materials for "Transcriptional differences for COVID-19 Disease Map genes between males and females indicate a different basal immunophenotype relevant to the disease"

**Supplementary Figure 1:** COVID-19 case fatality rates for males and females across 33 different countries or regions with over 1000 reported sex-disaggregated death data. The data is collected by Global Heath 50/50 on September, 17th, 2020.

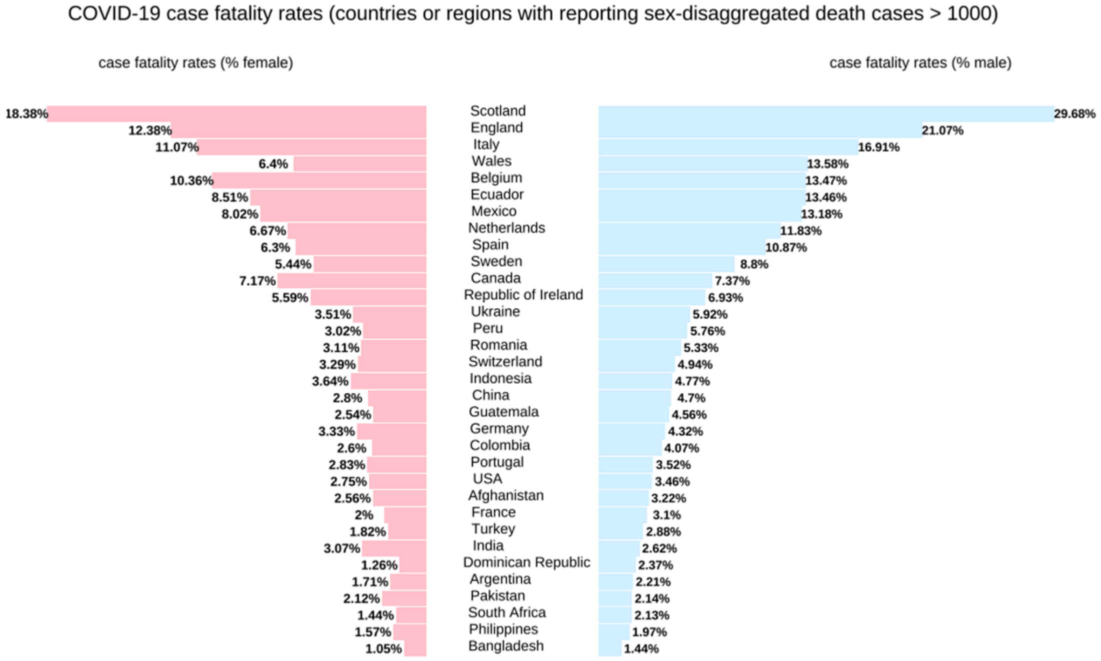

**Supplementary Figure 2:** The Principal Component Analysis plot for GTEX RNA-seq data of whole blood. Each dot corresponds to an individual. Data are colored by the cause of death. The PCA shows that the samples labelled as "ventilator case" strongly clustered apart from the rest of the samples in the first PC. These samples were therefore excluded from the DeCovid app.

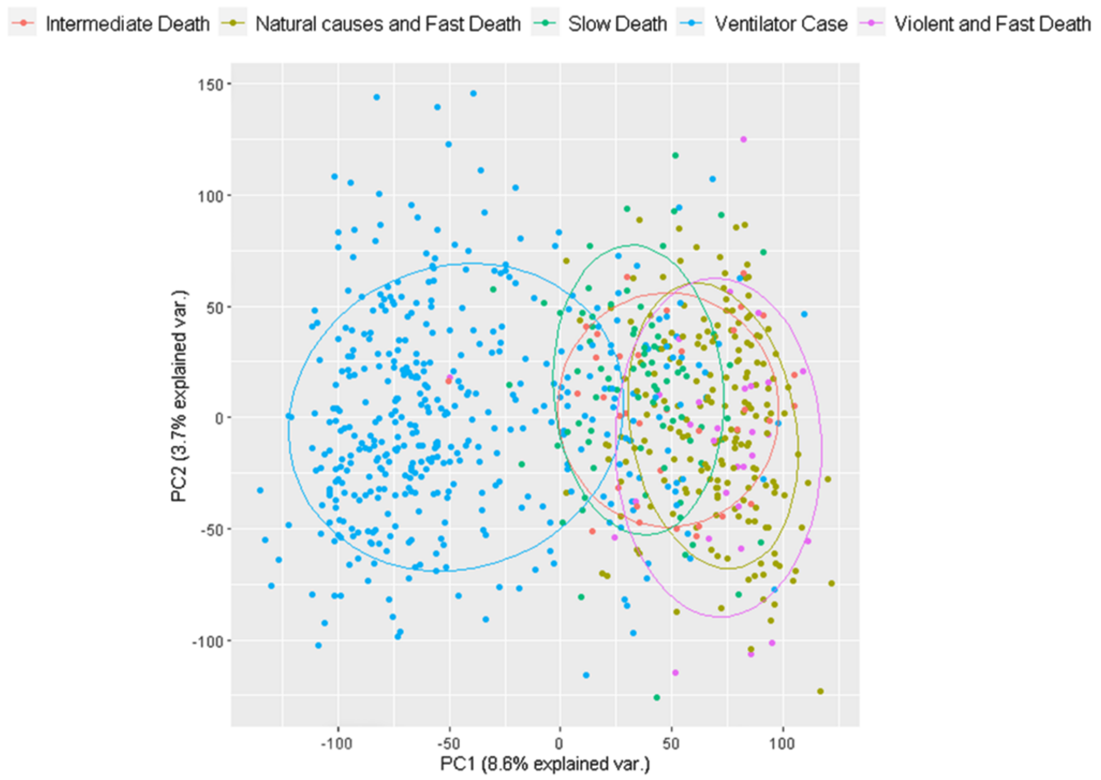
